# Anti-tau antibodies targeting a conformation-dependent epitope selectively bind seeds

**DOI:** 10.1101/2023.05.04.539475

**Authors:** Brian D. Hitt, Ankit Gupta, Ruhar Singh, Ting Yang, Joshua D. Beaver, Ping Shang, Charles L. White, Lukasz A. Joachimiak, Marc I. Diamond

**Author notes:** Corresponding Author: Marc Diamond, M.D., NS8.334, 6124 Harry Hines Blvd. Dallas, TX 75390. Equal contribution (Brian D. Hitt, Ankit Gupta).

## Abstract

Neurodegenerative tauopathies are caused by transition of tau protein from a monomer to a toxic aggregate. They include Alzheimer disease (AD), progressive supranuclear palsy (PSP), corticobasal degeneration (CBD), and Pick disease (PiD). We have previously proposed that tau monomer exists in two conformational ensembles: an inert form (M_i_), which does not self-assemble; and seed-competent form (M_s_), which self-assembles and templates ordered assembly growth. We proposed that cis/trans isomerization of tau at P301, the site of dominant disease-associated S/L mis-sense mutations, might underlie the transition of wild-type tau to a seed-competent state. Consequently, we created monoclonal antibodies using non-natural antigens consisting of fluorinated proline (P*) at the analogous P270 in R1, biased towards the trans-configuration at either the R1/R2 (TENLKHQP*GGGKVQIINKK) or R1/R3 (TENLKHQP*GGGKVQIVYK) interfaces. Two antibodies, MD2.2 and MD3.1 efficiently immunoprecipitated soluble seeds from AD and PSP, but not CBD or PiD. They stained brain samples of AD, PSP, and PiD, but not CBD. They did not immunoprecipitate, or immunostain tau from control brain. Creation of potent anti-seed antibodies based on the trans-proline epitope implicates local unfolding around P301 in pathogenesis. MD2.2 and MD3.1 may also be useful for therapy and diagnosis.

**Summary:** Tau protein undergoes conformational change to self-assemble and trigger neurodegeneration. We have proposed local unfolding events centered on P301 cis/trans isomerization, which expose amyloidogenic sequences. We used a non-natural peptide with a trans-proline to generate monoclonal antibodies that distinguish tau seeds from native tau in human brain. In addition to being important therapeutic and diagnostic leads, the activity of these antibodies supports structural studies implicating local conformational change in tau that underlies disease initiation.

## Introduction

Deposition of tau in ordered assemblies underlies Alzheimer disease (AD) and related neurodegenerative tauopathies, including progressive supranuclear palsy (PSP), corticobasal degeneration (CBD), and Pick disease (PiD)(Lee et al., 2001). These disorders are associated with distinct insoluble tau assembly structures that have been resolved by cryogenic electron microscopy (cryo-EM)(Scheres et al., 2020). Considerable experimental data supports the hypothesis that tau pathology progresses via trans-cellular propagation of protein aggregates, in which tau assemblies form in one cell, are released into the extracellular space, and are taken up by neighboring cells, where they act as templates to convert native tau into a growing aggregate (Gibbons et al., 2019; Goedert et al., 2017). This process, termed “seeding,” is readily replicated in simple cell models based on expression of full-length tau (Frost et al., 2009) or tau repeat domain (RD) (Holmes et al., 2014) fused to fluorescent proteins. When complementary fluorescent protein fusions are co-expressed in “biosensor” cells, aggregation is easily detected by fluorescence resonance energy transfer (FRET) (Hitt et al., 2021; Holmes et al., 2014). Biosensor cells are useful to characterize tau seeding activity in mouse models and human tissues, and detect tau seeds prior to development of neurofibrillary tau pathology in humans (Furman et al., 2017; Stopschinski et al., 2021), and mice (Holmes et al., 2014; Mirbaha et al., 2022).

The tau RD (amino acids 244-378) forms the backbone of amyloid cores in pathological tau deposits (Crowther et al., 1989; Fitzpatrick et al., 2017). Distinct conformations of the RD underlie the seeding and strain identity of pathological tau (Mirbaha et al., 2018; Sharma et al., 2018). Strains, by definition, propagate faithfully *in vivo* and cause distinct patterns of pathology. We have previously propagated tau strains derived from distinct tauopathies in cultured biosensor cells. These create unique patterns of neuropathology, rates of progression, and regional involvement upon inoculation into a transgenic mouse model (Kaufman et al., 2016; Sanders et al., 2014; Vaquer-Alicea et al., 2021). Others, by inoculating insoluble material from tauopathy brains, have also produced different patterns of pathology in mouse models (Boluda et al., 2015; Clavaguera et al., 2013; Clavaguera et al., 2009). Subsequently, cryo-EM has described in atomic detail tau assemblies from different tauopathies and documented distinct conformations of the RD (Shi et al., 2021). The simplest interpretation of these data is that individual tauopathies are caused by unique, disease-specific tau assemblies(Vaquer-Alicea et al., 2021).

The mechanistic origins of tauopathy are mysterious and likely diverse. We have previously concluded that tau monomer exists in two general conformational ensembles, an inert form (M_i_) that does not readily self-assemble or act as a template, and a seed-competent form (M_s_) that can self-assemble or trigger further refolding and assembly of monomers into oligomers (Mirbaha et al., 2018). We have described methods to produce M_s_ *in vitro* (Hou et al., 2021). M_s_ has myriad sub-structures, as isomers isolated from human brain or cell lines or encode distinct strains (Sharma et al., 2018). M_s_ is the earliest form of seed-competent tau we have detected in a mouse model, preceding formation of larger assemblies (Mirbaha et al., 2022), and we have linked formation of certain forms of M_s_ to changes in local tau structure (Hou et al., 2021; Mirbaha et al., 2018). Specifically, based on cross-linking mass spectrometry, molecular dynamic modeling, and biochemical studies, we have proposed that cis-trans proline isomerization at P301 underlies formation of M_s_ in certain cases (Chen et al., 2019; Drombosky et al., 2018; Mirbaha et al., 2018), with amyloid-forming motifs VQIINK and VQIVYK relatively more exposed in seed-competent (M_s_) vs. inert (M_i_) tau. We have observed that a trans-proline isomer at residue 270/301 (depending on 3R vs. 4R tau) favors an “open” conformation *in vitro* that is predicted to expose these amyloidogenic motifs (Chen et al., 2019).

To test these ideas further, we have now used linear peptide antigens with a synthetic fluorinated trans-proline residue to produce monoclonal antibodies against this region of tau (Figure 1). We employed a novel screening and selection protocol to enrich for seed specificity. We selected two novel monoclonal antibodies produced by this method, MD2.2 and MD3.1, and have characterized their binding characteristics *in vitro* with recombinant protein, brain lysates, and immunohistochemistry.

**Figure 1:**
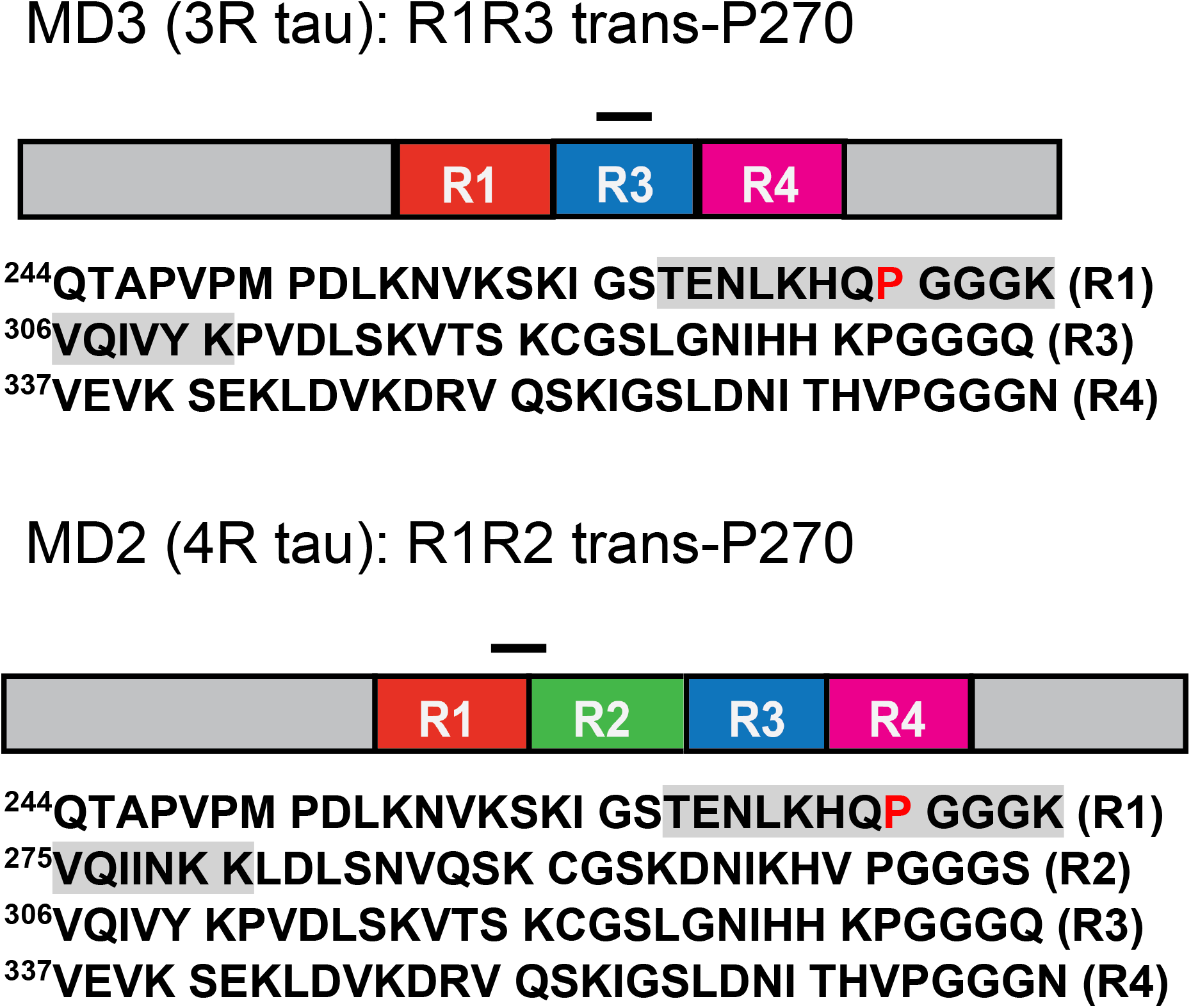
Peptide antigens for the MD antibody series. The MD2.2 and MD3.1 epitopes (grey) comprise amino acids 263-281 (4R) and 263-311Δ275-305 (3R) of the tau sequence, respectively. Both include a fluorinated proline (N-Boc-trans-4-fluoro-L-proline) at residue 270, shifting the equilibrium in favor of the trans-proline conformation.

## Materials and Methods

### Brain tissue

We identified cases from the brain bank of the Alzheimer’s Disease Center at UT Southwestern, selecting those with a single histopathological diagnosis of either AD, PSP, CBD, or PiD based on standard neuropathological evaluation, and excluding mixed pathologies. Negative control brains derived from cases multiple system atrophy due to α-synuclein accumulation, and frontotemporal lobar degeneration due to TDP-43 accumulation. We chose eight cases per disease for formalin-fixed, paraffin-embedded (FFPE) tissue and four cases per disease for frozen tissue. We included frozen frontal cortex from three healthy control brains, free of any detectable neuropathology. We used cores of middle frontal gyrus from each formalin fixed brain for tissue microarray (TMA) construction (below), and frozen samples of frontal gyri for soluble protein brain extracts. We performed a detergent-free homogenization of all frozen brain samples in 10% w/vol of 1x TBS with protease inhibitor cocktail (Roche) by mechanical disruption at 4°C followed by intermittent water bath sonication (Qsonica) for 5 minutes total “on” time. We centrifuged homogenates at 21,000 x g for 15 minutes at 4°C to remove cellular debris and determined protein concentrations by BCA assay (ThermoFisher).

### Generation of MD series of antibodies

We produced a series of novel monoclonal anti-tau antibodies (MD series) with a contract research organization (CRO, Genscript). The CRO synthesized linear peptides with sequences corresponding to amino acids 263-281 (R1R2) and 263-311Δ275-305 (R1R3) of tau (assuming 2N4R, 441 aa) for the MD2 and MD3 antigens, respectively, substituting N-Boc-trans-4-fluoro-L-proline at residue 270, and conjugated them to keyhole limpet hemocyanin (KLH). The CRO provided antisera from inoculated mice for screening with immunoprecipitation (IP)-seeding assay. We incubated supersaturating concentrations of antisera with Dynabeads Protein A (ThermoFisher) and used them in immunoprecipitation (IP) of tau from 10μg total soluble protein from an AD brain. We eluted protein in low pH elution buffer (Pierce) equivalent to starting IP/supernatant volume and neutralized with 1:10 1M Tris pH 8.5. We measured the tau seeding activity in equivalent volumes of IP eluent and supernatant with a cell-based tau seeding assay (below). We identified the mice with antisera most efficient in immunoprecipitating AD tau seeds and the CRO used these mice for thymus cell fusion to produce hybridoma clones. We analyzed supernatants from parental clones and subsequently subclones as above, including IP of seeds from non-AD tauopathies, and selected two hybridomas for detailed analysis: MD2.2 and MD3.1.

### Surface interferometry

We measured the kinetics of antibody association and dissociation using surface interferometry (Octet Red 384, FortéBio). To study antibody binding to peptides, we first loaded pre-equilibrated streptavidin biosensor probes with 400 nM of biotinylated peptides in 1X kinetics buffer (Sartorius) for 120 s, before a 180 s association and 180 s dissociation step. The peptides consisted of: R1/R2, R1/R2-transPro and R1/R3 and R1/R3-transPro. We next tested binding to antibodies by incubation with serially diluted preparations of HJ8.5, MD2.2, and MD3.1.

To measure binding to recombinant full-length (FL) tau monomer (tau 2N4R) and truncated tau monomer without the second repeat (tau 2N3R), we loaded the pre-equilibrated AMC biosensors (in 1X kinetics buffer from Sartorius) with 100 nM HJ8.5, MD2.2, and MD3.1 antibodies for 120 s, before a 120 s association step (except MD3.1) and 180 s dissociation step, with serial dilutions of 2N4R and 2N3R tau monomer proteins. MD3.1 showed a drift after 80 s of association (possibly due to the dissociation of immobilized antibody from biosensors), hence, we carried out the binding studies of MD3.1 with 80 s association and 180 s dissociation.

Finally, we used Octet System Data Analysis Software (FortéBio) to fit the kinetic curves to equations based on 1:1 kinetics and calculate K_D_, k_a_ (on), and k_d_ (off). We performed double referencing (sample and biosensor) for all our measurements to background subtract nonspecific binding.

### Immunoprecipitation

50 μl of Dynabeads Protein A (ThermoFisher) were washed per the manufacturer protocol and incubated with 10μg of monoclonal antibodies for 1 hour at room temperature. We then added washed beads to 10μg total soluble protein diluted in binding buffer (PBS + 0.02% Tween 20). We collected supernatants eluted captured proteins in low pH elution buffer (Pierce), and neutralized the buffer with 1:10 1M Tris pH 8.5 with a final volume equivalent to the supernatant.

### Western blotting

We performed SDS-PAGE on IP and supernatant samples equivalent to 2μg total starting protein with 4-12% Bis-Tris gels. We transferred protein to a PVDF membrane, blocked with 5% dry milk, and probed with a 1:2000 dilution of anti-tau rabbit polyclonal antibody, followed by a goat anti-rabbit secondary, and developed the blots with the ECL Ultra detection kit (ThermoFisher). For unprocessed gel images, please see Supplemental Figures 4,5.

### Tau seeding assay

We performed tau seeding assays using the tau RD P301S v2H HEK biosensor cell line as previously described(Hitt et al., 2021). For each brain we determined the optimal total mass to use for tau seeding assays using a serial dilution dose response curve to ensure the amount used was well within the linear range in each case. We incubated the equivalent volume of IP and supernatant with Lipofectamine 2000 transfection reagent and added them to 3 wells per sample of biosensor cells in a 96-well plate. We collected cells after 48 hours and fixed them in 2% PFA for 10 minutes before measuring % FRET by flow cytometry as previously described (Furman et al., 2015).

### Tissue microarrays

We constructed a tissue microarray (TMA) containing samples from each tauopathy case, including cores of FFPE middle frontal gyrus and cerebellum (as negative control) from eight brains per disease using an Epredia 3DHistech TMA Master II (Richard-Allan Scientific, Kalamazoo, MI) robotic tissue micorarrayer and 1.6 mm diameter cores. Donor core sites were determined from overlays of H&E-stained scout sections of each donor block, prepared on an Aperio CS2 (Leica Biosystems, Buffalo Grove, IL) whole slide imager. We mounted 5μm sections from the TMA block on glass slides and performed immunohistochemical staining via a Bond III (Leica) automated immunostaining platform with optimal dilutions of each primary antibody determined by an initial serial dilution to minimize background while retaining foreground staining in sections with pathology. Dilutions were 1:200 AT8 (ThermoFisher), 1:16000 MD2.2, 1:4000 MD3.1, 1:32000 HJ8.5. We captured high resolution images of whole slides with an optical scanner and selected qualitatively representative fields from each stained tissue core.

### Modeling of flanking sequences of amyloid fibrils

To model the full 2N4R tau sequence in the context of cryo-EM assemblies we employed a comparative procedure in Rosetta (Song et al., 2013) leveraging the AD, PSP, and CBD fibril conformations as templates (PDBids: 5O3L, 6TJO and 7P65). Fragments for full-length 2N4R tau were built using the fragment picker tool and 500 models of FL tau monomers were built for AD, PSP and CBD using the homology modeling module (Song et al., 2013). Individual monomers were assembled into trimers using structural alignment to the cryo-EM template and trimeric assemblies were scored. The lowest scoring models were subsequently minimized using constraints derived from the trimeric fibril core template. Next, to evaluate the exposure of the different epitopes (HJ8.5, residues 25-30; R1R2, residues 263-280), we calculated the per-residue solvent accessible surface area (SASA) using the SasaCalc class in PyRosetta version across the AD, PSP and CBD trimeric ensembles (Chaudhury et al., 2010). SASA was calculated with 1.4 Å probe radius. SASA distributions for the AD, PSP and CBD trimeric ensembles were calculated for the R1R2 (MD2.2), C-term of R1R2 (VQIINK) and HJ8.5 epitopes. The data were plotted with seaborn library and matplotlib in python3.7 modules. For the MD2.2 epitope, we also computed the SASA distributions for a fraction of the sequence to identify the minimal sequence that distinguishes PSP from CBD SASA in the MD2.2 epitope. For simplicity, we focused on the MD2.2 epitope in the context of 4R tau, which is present in all disease-associated assemblies.

## Results

### MD2.2 and MD3.1 each bind FL tau and peptide antigens

MD2.2 and MD3.1 were raised against 263-281 (R1R2) and 263-311Δ275-305 (R1R3) tau sequences, respectively, with fluorinated proline at residue 270 position (Figure 1). After their original selection based on binding AD seeds, we compared the binding characteristics of MD2.2 and MD3.1 to HJ8.5, a high affinity antibody that binds the N-terminus of tau (Yanamandra et al., 2013).

We first used surface interferometry to determine the affinity of HJ8.5, MD2.2 and MD3.1 for recombinant full-length (FL) tau (3R and 4R). HJ8.5 bound both tau isoforms with similar affinity in the low nanomolar range. MD2.2 and MD3.1 each bound 3R tau with slightly higher affinity than 4R tau, and with a K_D_ ∼5-10x greater than HJ8.5 (i.e. lower affinity) (Figure 2). We next checked binding to short peptides: R1R2, trans-P-R1R2, R1R3, and trans-P-R1R3. MD2.2 and MD3.1 each bound them with ∼10x higher affinity than FL protein (Figure 2B). Consistent with the antigens used to create them, MD2.2 and MD3.1 bound the trans-P-R1R2 and trans-P-R1R3 peptides with K_D_ 2-3-fold higher than for trans-P-R1R2 and trans-P-R1R3 peptides (Figure 2B, Supplemental Figure 1). HJ8.5, which binds the N-terminus of tau, did not bind any of the peptides (Supplemental Figure 1). Taken together, our data indicated MD2.2 and MD3.1 each had higher affinity for trans-P configurations, while still binding FL tau isoforms. Because there was no difference in 3R vs 4R tau binding, the antibodies apparently bound epitopes common to both isoforms.

**Figure 2:**
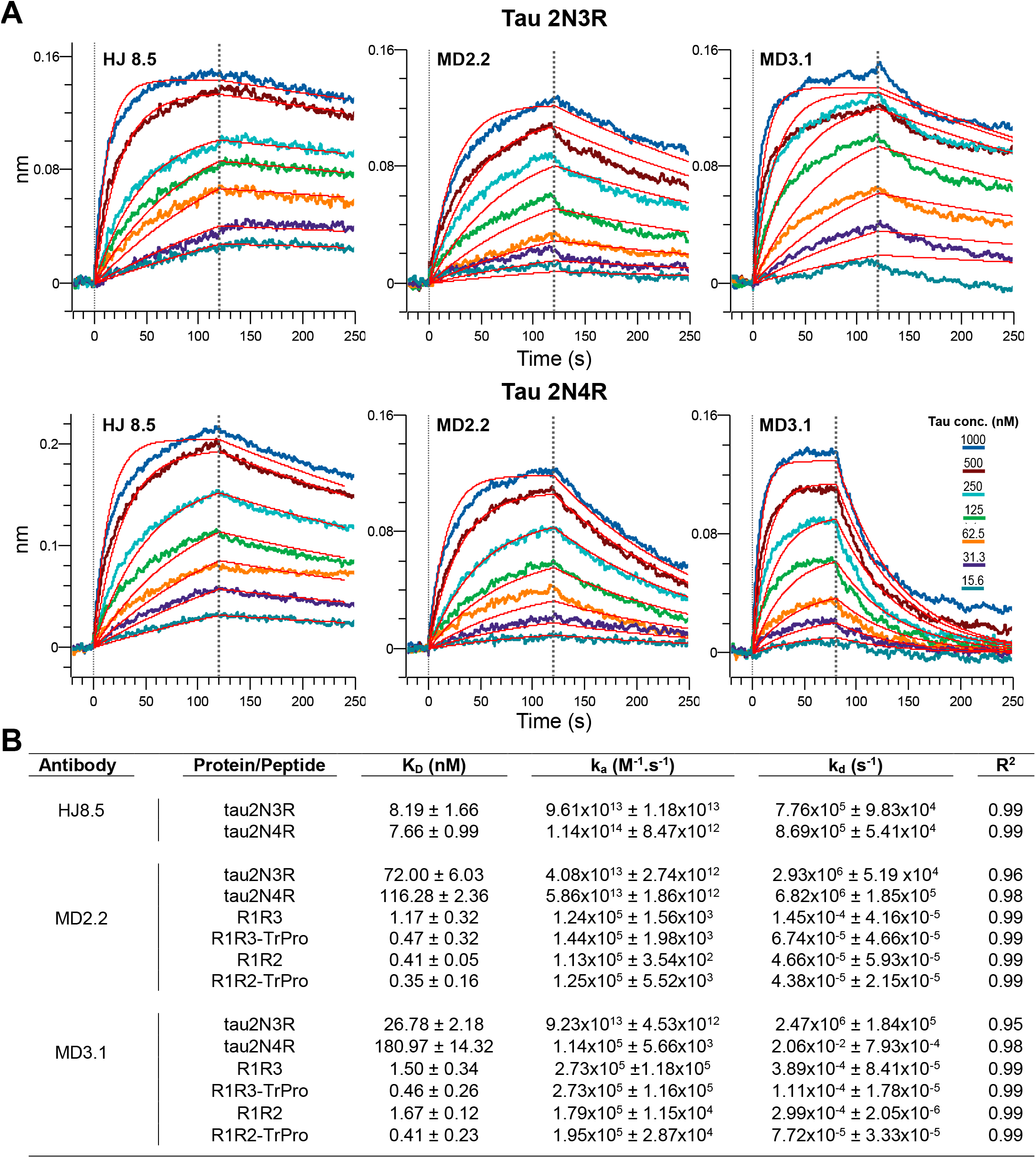
Binding properties to recombinant tau and peptide antigens. (**A**) Kinetic traces for full-length 2N3R (upper panel) and 2N4R (bottom panel) recombinant tau. The kinetic traces are fitted to 1:1 kinetics. For MD2.2 and MD3.1, which target the repeat domain containing multiple similar PGGG containing repeats, the curves show fast-on, fast-off kinetics with a poorer fit of 1:1 kinetics. (**B**) K_D_, k_a_, k_d_, and R^2^ for HJ8.5, MD2.2, and MD3.1 were calculated based on a grouped curve fit of 7 tau concentrations with 1:1 binding parameters. Data for HJ8.5 binding with various peptides is not shown (see Supplemental Figure 1).

### Immunoprecipitation of tau vs. tau seeds from AD vs. control brain

We next tested binding properties of MD2.2 and MD3.1 for seeds from control vs. AD brain. We prepared soluble homogenates of frozen frontal cortex from three healthy control and three AD brains. We used HJ8.5, MD2.2 and MD3.1 to immunoprecipitate (IP) samples. MD2.2 and MD3.1 did not bind detectable tau from control brains, whereas HJ8.5 bound nearly all of it (Figure 3A,C,E). When MD2.2 and MD3.1 were used to IP AD brain we detected very faint signal, whereas HJ8.5 immunoprecipitated most soluble protein. Using tau RD(P301S) v2H biosensor cells (Hitt et al., 2021) we detected no seeding activity in the control brain homogenates (Figure 3B, D, F). We detected robust seeding activity in all three AD brains, observed in the supernatant fractions of the control IPs using non-specific mouse IgG (Figure 3H,J,L). In contrast to the western blots, MD2.2 and MD3.1 immunoprecipitated virtually all detectable AD seeds (Figure 3H,J,L). This contrasted with HJ8.5, which precipitated nearly all total soluble tau, but left significant seeding activity in the supernatant in all cases. The MD2.2 and MD3.1 antibodies therefore bound a small subset of soluble tau that was absent from control brains and accounted for essentially all seed-competent tau in AD brains.

**Figure 3:**
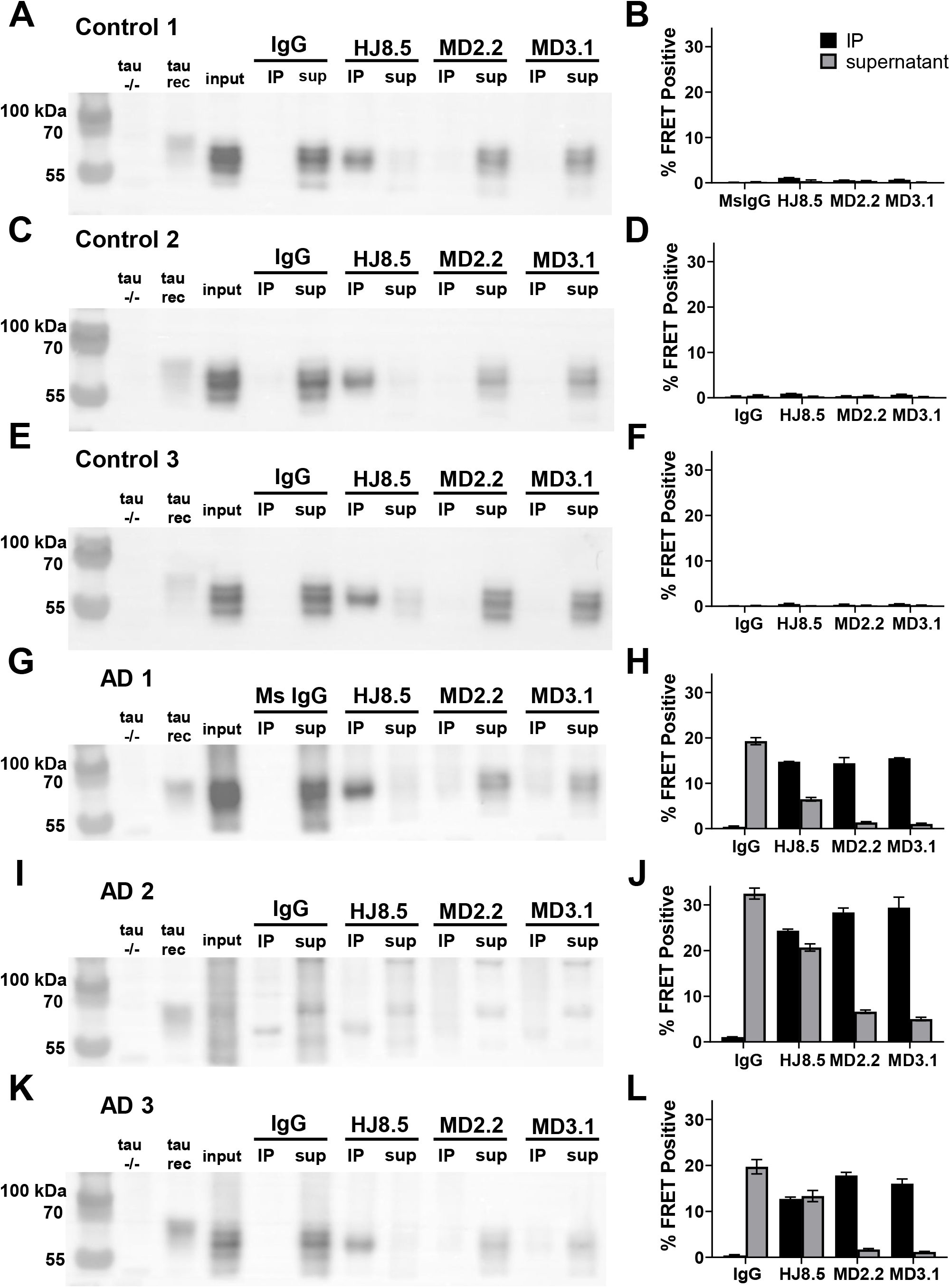
MD2.2 and MD3.1 bind soluble tau AD seeds. Western blots of total tau in the IP and supernatant fractions from immunoprecipitation of three control brains **(A, C, E)** and three AD brains **(G, I, K)** indicated that MD2.2 and MD3.1 bound no tau from control brains, and only a very small fraction of tau from AD brains. By contrast, HJ8.5 bound nearly all tau from all brains. Seeding assays of the IP and supernatant fractions showed that nearly all seeding activity from AD brains was in the IP fractions of MD2.2 and MD3.1, whereas much was left in the supernatant by HJ8.5 **(H, J, L)**. No significant seeding activity was detected in control brains **(B, D, F)**. Input was the pre-IP total protein for each brain; tau-/-negative control was the equivalent total protein from a tau-/-mouse brain, tau rec indicates recombinant tau monomer (2N4R). Non-specific mouse IgG was the negative control for immunoprecipitation.

### MD2.2 and MD3.1 discriminate seeds from different tauopathies

Tau assembly structures vary among tauopathies. To test the specificity of MD2.2 and MD3.1 for AD vs. other tauopathies, we prepared soluble homogenates of frozen frontal cortex from corticobasal degeneration (CBD), progressive supranuclear palsy (PSP), and Pick disease (PiD) brains. We detected no tau by WB in the IP fractions of MD2.2 and MD3.1, whereas HJ8.5 immunoprecipitated virtually all detectable tau (Figure 4A,C,E). All three tauopathy brains exhibited tau seeding in v2H biosensors (Figure 4B,D,F). MD2.2 failed to efficiently IP virtually any seeding activity from CBD and PiD brains, while binding PSP seeds. MD3.1 also bound seeds well from the PSP brain, but with much lower efficiency for CBD and PiD. HJ8.5 failed to IP CBD brain efficiently, but did so for PiD and PSP. In summary MD2.2 and MD3.1 efficiently bound tau associated seeds from AD and PSP, but not from CBD and PiD.

**Figure 4:**
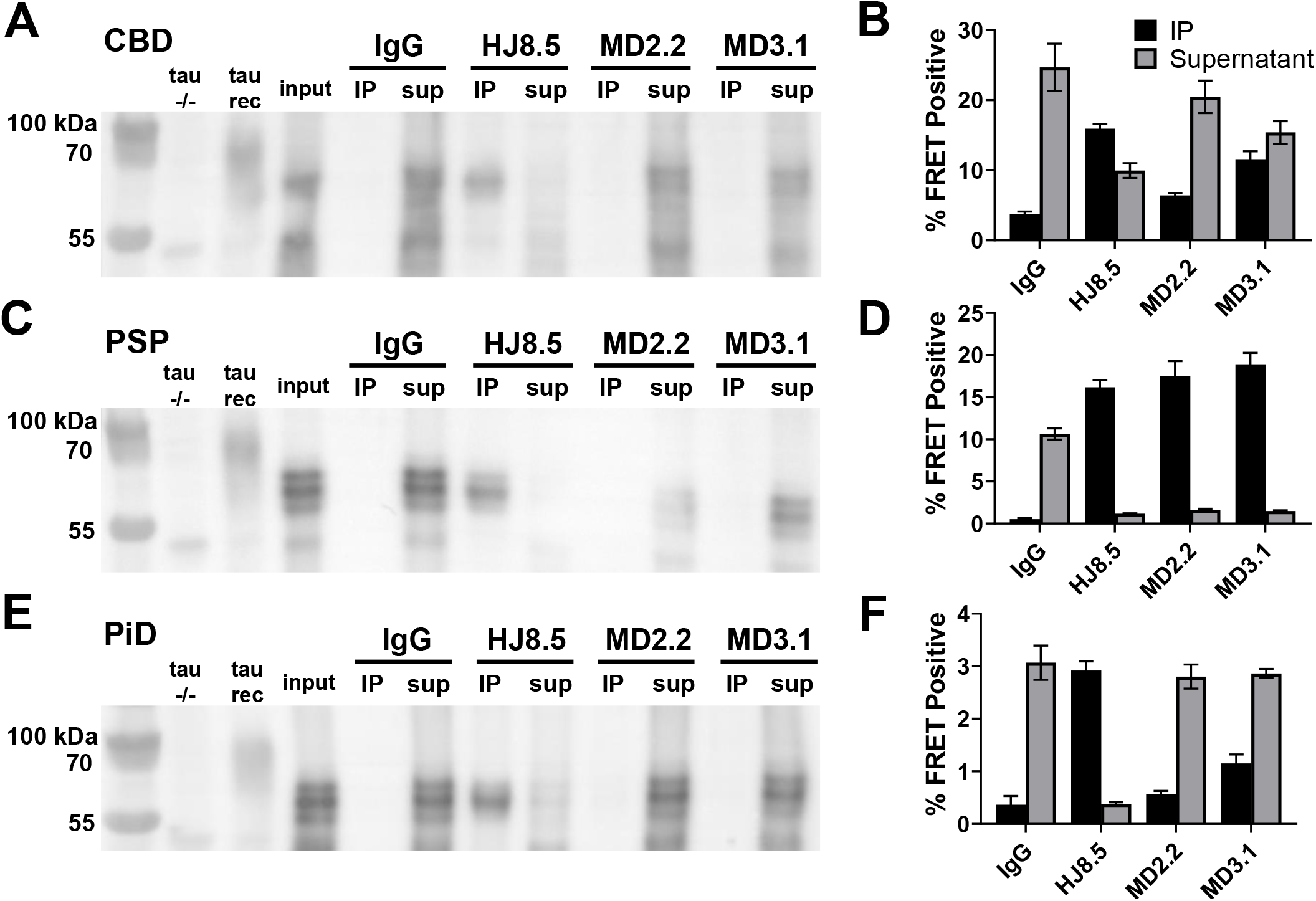
MD2.2 and MD3.1 exhibit specificity for tauopathy seeds. IP of brain protein from the non-AD tauopathies CBD **(A)**, PSP **(C)**, and Pick disease **(E)** demonstrated undetectable total tau by MD2.2 and MD3.1 vs. most tau vs. HJ8.5, which precipitated most tau. Seeding assays revealed significant seeds from CBD **(B)** and PiD **(D)** in the supernatant after IP with MD2.2 and MD3.1. Conversely, nearly all PSP seeds were bound by MD2.2 and MD3.1 **(D)**. Input was the pre-IP total protein for each brain; tau-/-negative control was the equivalent total protein from a tau-/-mouse brain, tau rec indicates recombinant tau monomer (2N4R). Non-specific mouse IgG was the negative control for immunoprecipitation.

### MD2.2 and MD3.1 discriminate tauopathies *in situ*

We stained adjacent sections prepared from TMA blocks with MD2.2, MD3.1, HJ8.5, and AT8, the standard phospho-tau antibody used to identify pathological tau (Goedert et al., 1995). MD2.2 and MD3.1 stained tau in AD, PSP, and PiD but not CBD or brains containing α-synuclein or TDP-43 pathology (Figures 5,6,7). In AD, MD2.2 and MD3.1 preferentially stained perinuclear and cell body tau inclusions (Figure 5A), particularly in brains with lighter AT8 staining. In contrast, AT8 exhibited more diffuse staining throughout the neuropil, and HJ8.5 featured high background staining with poor signal-to-noise (Figure 5A). In PSP brains, MD2.2 and MD3.1 stained only small, granular perinuclear rings of accumulated tau, and not tangles or tufted astrocytes like AT8 (Figure 5B). Overall the staining patterns for MD2.2 and MD3.1 revealed a small fraction of total AT8 signal in AD and an extremely small fraction in PSP. In CBD brain sections we observed no staining by either antibody (Figure 6A). In PiD sections MD2.2 and MD3.1 exhibited some Pick body labeling, albeit much less than either AT8 or HJ8.5 (Figure 6B). Neither MD3.1 or MD2.2 stained other neurodegenerative disease brain samples without tau pathology, an MSA brain featuring α-synuclein aggregates, or an FTLD brain featuring TDP-43 aggregates (Figure 7). Taken together, the staining characteristics of MD2.2 and MD3.1 reflected specificity for different tau assembly structures, as distinct from AT8.

**Figure 5:**
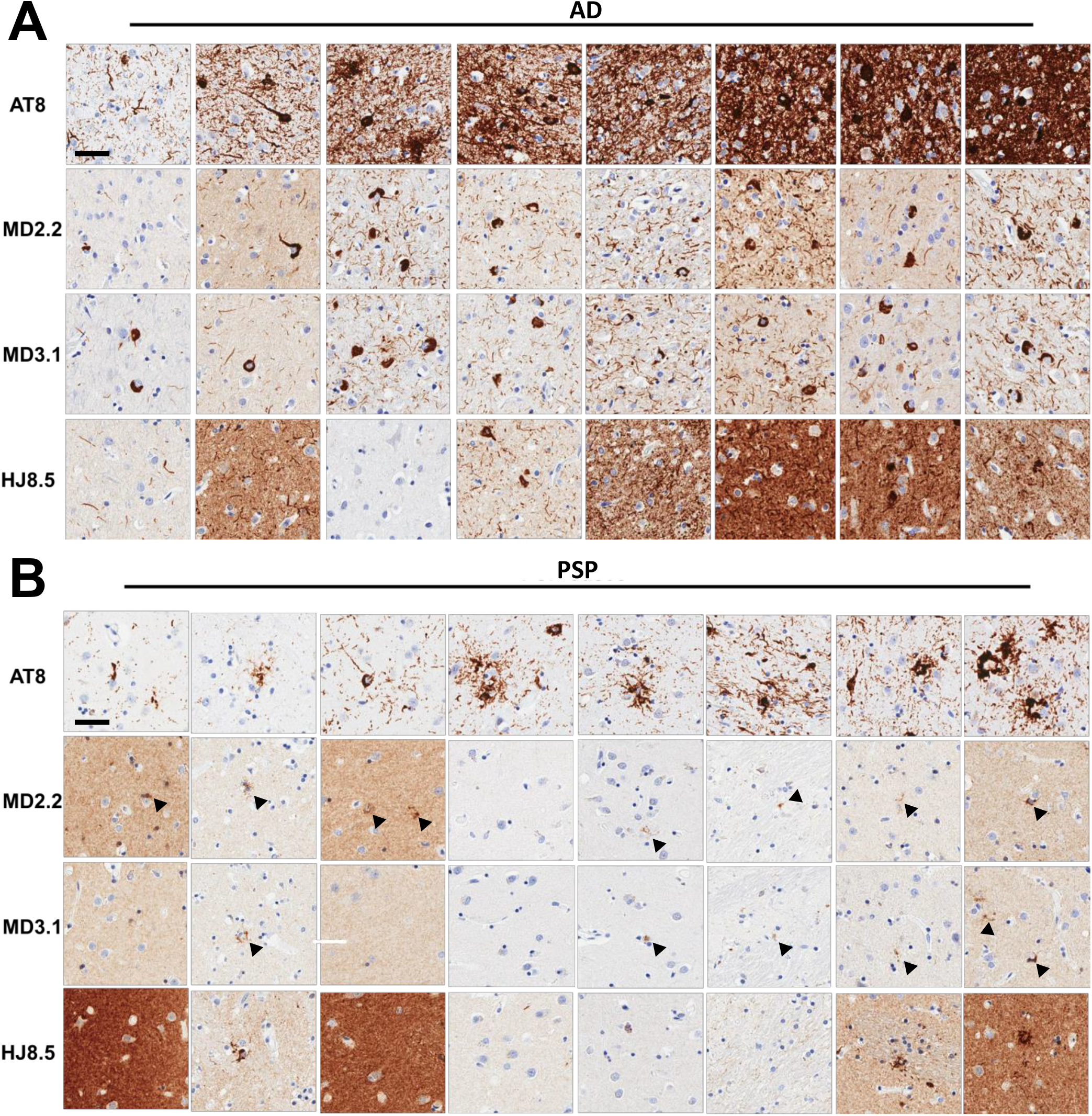
MD2.2 and MD3.1 staining of AD and PSP brains. Representative images of tissue microarray IHC staining of fixed frontal cortex of eight AD brains **(A)** and eight PSP brains **(B)** showed less total tau staining on average by MD2.2 and MD3.1 than by HJ8.5, and a distinct pattern of tau aggregates vs. AT8, with perinuclear granular bodies noted particularly in PSP (arrowheads). Scale bar = 50 μm.

**Figure 6:**
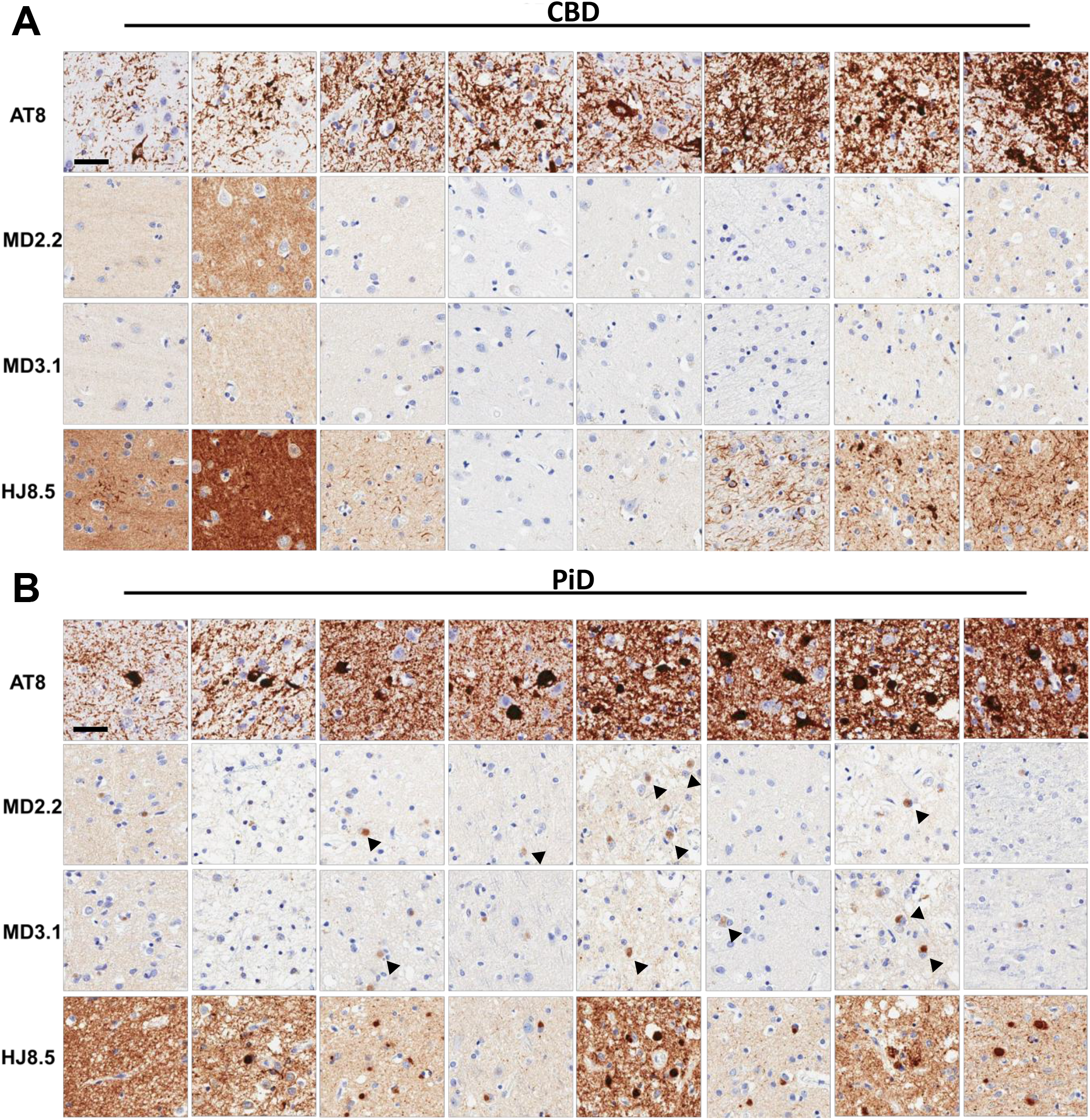
MD2.2 and MD3.1 staining of CBD and PiD brains. Representative images of tissue microarray IHC staining of fixed frontal cortex of eight CBD brains **(A)** and eight Pick disease brains **(B)**. CBD brain exhibited no significant tau staining above background. In PiD there was faint labeling of Pick bodies (arrowheads). This contrasted with staining by AT8 and HJ8.5. Scale bar = 50 μm.

**Figure 7:**
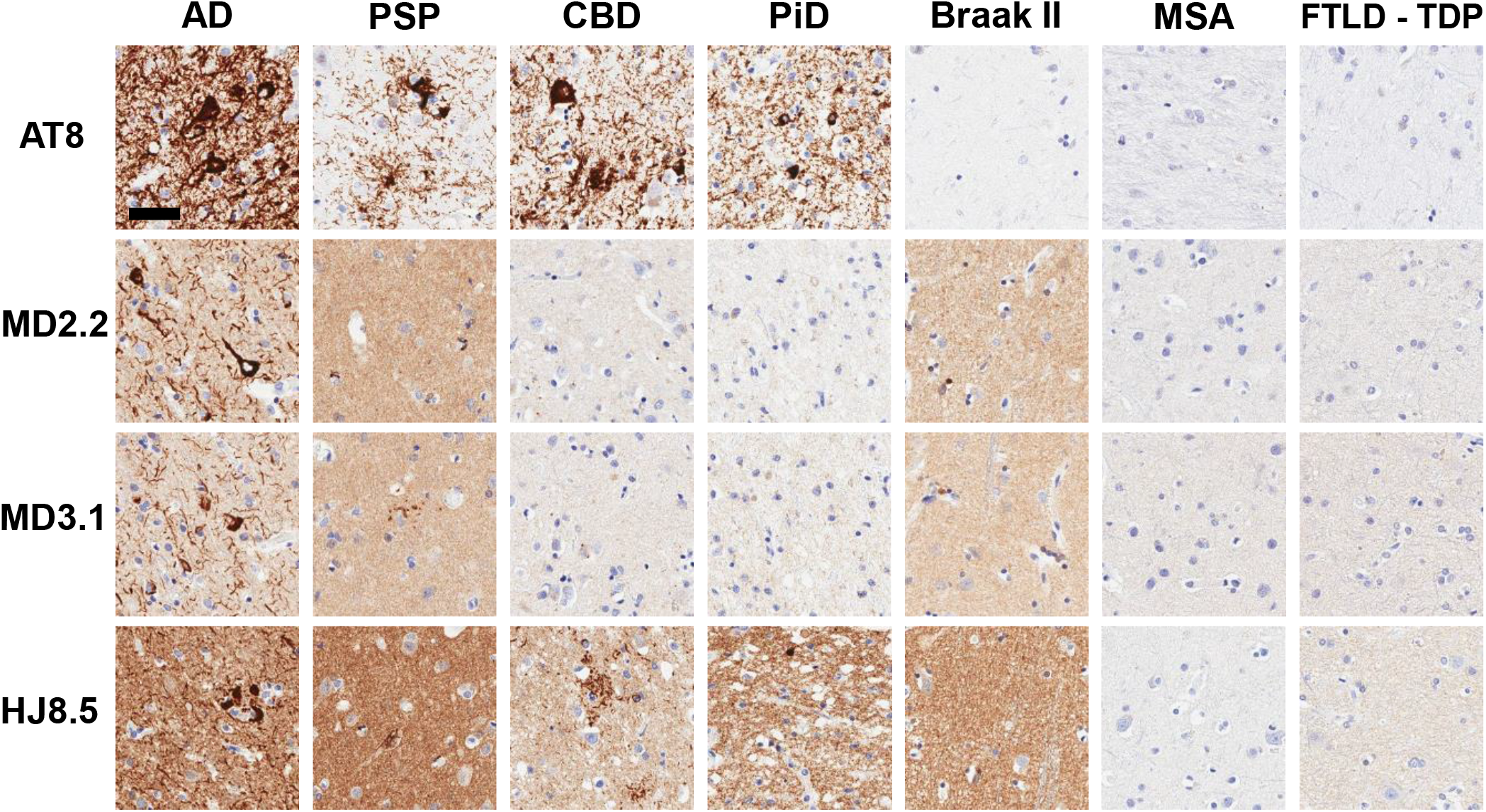
MD2.2 and MD3.1 staining of AD and PSP vs non-tauopathy brains. Immunohistochemistry of brains with AT8(+) pathology indicated more MD2.2 and MD3.1 staining in AD and PSP than in CBD and PiD. The Braak II sections are from a Lewy body dementia brain with AD NFTs only in limbic tissue, and exhibited staining with MD2.2 and MD3.1. In non-tauopathy MSA or FTLD-TDP brains there was no staining. Scale bar = 50 μm.

## Discussion

This study tested a prediction based on earlier work which indicated that tau monomer exists in distinct conformational ensembles, M_i_ vs M_s_, discriminated by local folding patterns which differentially expose motifs required for self-assembly. Specifically, we have proposed that M_s_, a trans-proline configuration of P301 unfolds a local hairpin to expose the critical aggregation motifs VQIINK or VQIVYK (Mirbaha et al., 2018). To test this idea, we used an immunogen with a non-natural trans-proline residue to vaccinate mice and derived two monoclonal antibodies: MD2.2, directed at a trans-P-R2R3 epitope (4R); and MD3.1, directed at a trans-P-R1R3 epitope (3R). These antibodies had noteworthy characteristics. Both bound recombinant FL tau monomer *in vitro*, based on surface interferometry. However, when used to IP tau derived from control brains they did not bind detectable tau, and with tauopathy brains they precipitated minimal detectable tau vs. a control N-terminal antibody (HJ8.5). They efficiently depleted seeds from AD and PSP lysates in a biosensor assay, while they bound those from CBD or PiD less efficiently. Both antibodies efficiently stained AD brain, but not CBD, with less efficient staining of PiD and PSP. Neither antibody stained control brain. Taken together with earlier studies, these data are consistent with the idea that local folding exposes unique epitopes in tau that enable aggregation, but that these events are disease (or strain)-specific.

### Differential binding based on strain-dependent epitope exposure

MD2.2 and MD3.1 are strain-specific, efficiently binding seeds of AD and PSP but not CBD or PiD. This could reflect distinct proteolytic processing or post-translational modifications. Alternatively, we hypothesized that the epitopes were differentially exposed. Cryo-EM has demonstrated distinct conformations of the amyloid cores from these four tauopathies, composed of beta sheets either two (AD, PiD), three (CBD), or four (PSP) layers (Shi et al., 2021). Consequently, we used published cryo-EM structures coupled with molecular dynamic modeling (Rosetta) of the unresolved epitopes to explain these binding characteristics. With an *ab initio* protocol to build trimers of the fibril cores, we modeled the flanking (“fuzzy coat”) regions of the N- and C-termini to include the full 2N4R tau sequence (Supplemental Figure 2A,B). We then computed solvent accessibly surface area (SASA) for the R1R2 epitope (residues 263-280) in each published disease-associated structure, with HJ8.5 as a control (Supplemental Figure 2C). Unsurprisingly, the model predicted the HJ8.5 epitope, far removed from the fibril cores, to be highly accessible. This was consistent with its ability to bind both tau monomer and assemblies. For the AD assembly, the R1R2 epitope was predicted to be highly exposed, possibly explaining the efficacy of antibodies in binding AD seeds. By contrast, the R1R2 epitope, and VQIINK amyloid motif, were predicted to be more buried in CBD and PiD. In PSP, the modeling predicted higher solvent accessibility of the epitope, although not as much as in AD (Supplemental Figure 2C, Supplemental Figure 3). Although it is not possible to structurally resolve highly mobile flanking sequence structure(s) within fibrils, the modeling and IP/seeding data were consistent with the idea that epitope accessibility might determine antibody avidity, at least for AD seeds vs. others.

### Soluble seed-competent tau is a small component of tauopathy brain tau

The differential binding of MD2.2 and MD3.1 to seed-competent vs. inert tau in brain revealed relative amounts of these species. While the antibodies had relatively high affinity for recombinant tau monomer *in vitro*, they failed to immunoprecipitate tau monomer from control brain. We hypothesized that the inert recombinant protein (M_i_) was relatively flexible, and might transiently adopt a diversity of conformations; that M_i_ within control brain was more rigidly folded or otherwise restricted to conformations that mask epitope accessibility; or a combination of both. The differential seed binding was striking, with virtually all seeding activity immunoprecipitated from AD and PSP brain lysates, but not CBD or PSP. Even with complete precipitation of seeds, only minimal tau monomer was detected on western blot (and no evidence of higher-order species). Thus, compared to total tau (as detected by HJ8.5 immunoprecipitation), seed-competent tau (whether monomer or assembly) appeared to represent tiny fraction of the soluble protein. This may explain why antibodies that bind all forms of tau, such as HJ8.5, failed to efficiently reduce pathology after peripheral administration—most tau bound would represent non-pathological species.

### Diagnostics and Therapeutics

Anti-tau antibodies have enabled pathological characterization of tauopathies, and anti-tau immunotherapy is a promising therapeutic strategy (Colin et al., 2020; Novak et al., 2018). Four anti-tau monoclonal antibody therapies, gosuranemab, tilavonemab, semorinemab, and zagotenemab, target epitopes N-terminal to the RD and have all failed in phase II clinical trials for either AD or PSP (Ayalon et al., 2021; Boxer et al., 2019; Dam et al., 2021; Hoglinger et al., 2021; Mullard, 2021a; Mullard, 2021b; Vaz and Silvestre, 2020). We note in this study that HJ8.5, the mouse version of tilavonemab, failed to efficiently IP all tau seeds compared to MD3.1 and MD2.2, despite efficient binding of full-length tau. As tau seeds likely must include the repeat domain, it is possible that proteolytic processing could remove N-terminal or C-terminal epitopes (Binder et al., 2005; Blennow et al., 2020; Cicognola et al., 2019; Sato et al., 2018), rendering an N-terminal antibody less effective.

We note that a humanized monoclonal antibody, E2814, also targets a repeat domain epitope (HVPPGG) present in both R2 and R4, and reduced *in vivo* tau deposition (Roberts et al., 2020). The epitope is fairly similar to the HQPGG sequence in R1 that was contained in the antigens used to create MD2.2 and MD3.1. The active immunotherapy AADvac1 also used an epitope similar to those for MD2.2 and MD3.1, but with a proline deletion rather than a synthetic trans-proline. This could have a similar effect on the secondary structure of the antigen. Importantly, despite multiple early anti-tau vaccine failures, upcoming clinical trials will answer questions about the efficacy and clinical utility of targeting conformation-specific RD epitopes. In addition to their potential therapeutic value, the specificity of these antibodies for tau seeds may facilitate their use as diagnostics to detect pathological species in biofluids.

## Figure Legends

**Supplemental Figure 1: BLI kinetics curves of antibody binding to various R1R2/R1R3 tau peptides**. (**A**) Kinetic traces for R1R2 (upper panel) and R1R2 trans-proline (bottom panel) peptides. The kinetic traces are fitted to 1:1 kinetics. (**B**) (A) Kinetic traces for R1R3 (upper panel) and R1R3 trans-proline (bottom panel) peptides. The kinetic traces are fitted to 1:1 kinetics. Whereas both MD2.2 and MD3.1 bound peptides, we observed no significant binding with HJ8.5.

**Supplemental Figure 2: Modelling of tau fibrils in AD, PSP, and CBD**. Rosetta was used to create ab initio models of published core structures within the context of FL tau to study the “fuzzy coat.” (**A**) Schematic illustration of model generation. (**B**) Schematic representation of full-length tau and repeat domain of tau highlighting the binding epitopes for HJ8.5, MD2.2, and MD3.1 antibodies. The repeat domains are colored red (R1), green (R2), blue (R3) and magenta (R4). The N- and C-termini of tau are colored grey. (**C**) Distinct ensemble representation of antibody binding regions R1R2 (MD2.2), C-term of R1R2 (VQIINK) and HJ8.5 using SASA values. AD, CBD and PSP SASA distributions are colored purple, yellow and cyan, respectively. Note the strong SASA score for AD fibrils bound by both MD2.2 and MD3.1. PSP exhibits a small degree of predicted accessibility vs. CBD. HJ8.5 is predicted to bind all structures equally.

**Supplemental Figure 3:** Energy distribution plot of assembled AD, CBD and PSP models in fibril form. Energy distribution of ensembles produced for AD, CBD and PSP trimers containing full length 2N4R tau. The energies are shown as Rosetta Energy Units (REU).

**Supplemental Figure 4: Unprocessed images of western blots used in Figure 3**.

**Supplemental Figure 5: Unprocessed images of western blots used in Figure 4**.

